# The evolutionary dynamics that retain long neutral genomic sequences in face of indel deletion bias: a model and its application to human introns

**DOI:** 10.1101/2022.07.25.501396

**Authors:** Gil Loewenthal, Elya Wygoda, Natan Nagar, Lior Glick, Itay Mayrose, Tal Pupko

## Abstract

Insertions and deletions (indels) of short DNA segments are common evolutionary events. Numerous studies showed that deletions occur more often than insertions in both prokaryotes and eukaryotes. It raises the question why neutral sequences are not eradicated from the genome. We suggest that this is due to a phenomenon we term *border-induced selection*. Accordingly, a neutral sequence is bordered between conserved regions. Deletions occurring near the borders occasionally protrude to the conserved region and are thereby subject to strong purifying selection. Thus, for short neutral sequences, an insertion bias is expected. Here, we develop a set of increasingly complex models of indel-dynamics that incorporate border-induced selection. Furthermore, we show that short conserved sequences within the neutrally evolving sequence help explain: (1) the presence of very long sequences; (2) the high variance of sequence lengths; (3) the possible emergence of multimodality in sequence length distributions. Finally, we fitted our models to the human intron length distribution, as introns are thought to be mostly neutral and bordered by conserved exons. We show that when accounting for the occurrence of short conserved sequences within introns, we reproduce the main features, including the presence of long introns and the multimodality of intron distribution.

## Introduction

Insertions and deletions (indels) of short DNA segments are common molecular evolutionary events (Cartwright, 2009), whose effect expands to macro-evolutionary processes, such as the divergence among species (Anzai et al. 2003; Britten 2002; Britten et al. 2003; Wetterbom et al. 2006). By analyzing homologous genomic sequences across various prokaryotic and eukaryotic taxa, it was repeatedly shown that deletions are more common than insertions (Fitch, 1973; Graur et al., 1989; De Jong and Rydén, 1981; Kuo and Ochman, 2009; Loewenthal et al., 2021; Mira et al., 2001; Ogata et al., 1996; Van Passel et al., 2007; Tao et al., 2008; Zhang and Gerstein, 2003), a phenomenon termed “deletion bias”. The deletion bias raises a question: why genomes and non-coding regions such as introns do not shrink over the course of evolution? Intriguingly, the opposite has supposedly happened, as eukaryotes have larger genomes (Cavalier-Smith, 1982), longer proteins (Brocchieri and Karlin, 2005), and much larger intergenic regions (Ahnert et al., 2008) compared to prokaryotes. Petrov (2002) suggested that the genome size is determined by two competing forces: short indels that reduce the genome size and large insertions (e.g., segmental duplications and the addition of transposable elements) that increase it. While this description may partially explain the overall genome size, it does not explain the length distribution of neutral sequences, such as introns, and the presence of short introns over long evolutionary time.

He et al. (2019) developed a deterministic model that describes how the length of a neutral sequence evolves given insertion and deletion rates, and assuming that only indels of length one are allowed. As one would expect, the sequence length grows exponentially when there is an insertion bias, but when the insertion and deletion rates are equal, the sequence length grows linearly. This result is quite counter-intuitive, and the authors explain it by noting that insertions emerge in between nucleotides and, thus, given a sequence of length *N* there are *N*+1 possible positions for insertions and only *N* possible positions for deletions. Under the setting of deletion bias, He et al. (2019) suggested that neutral sequences will be eliminated. However, the more elaborated statistical model TKF91 (Thorne et al., 1991) that similarly allowed for indels of size one only, demonstrated that under very weak deletion bias, neutral sequences will be maintained.

Indel dynamics may partially explain the distribution of intron lengths within and among organisms, and the length difference between introns of closely related species are correlated to indels (Moriyama et al., 1998; Ogata et al., 1996). Introns are non-coding sequences that are mostly neutral (Resch et al., 2007), but reside between exons, which are usually highly conserved (Siepel and Haussler, 2004). The distribution of the intron lengths is highly dispersed and thus it is usually plotted on a log scale. On such a scale it is often multimodal (Gotoh, 2018). For example, the distribution of human intron lengths is bimodal and ranges from 30 to 1,160,411 base-pairs (Piovesan et al., 2015). Introns length distributions of various organisms were fitted statistically with a Frechet mixture model, and demonstrated that in almost all eukaryotes, the log intron-length distribution is composed of multiple distinct components. This phenomenon was hypothesized to stem from the presence of alternative splicing mechanisms (Gotoh, 2018). Other studies classified introns according to their lengths and suggested that different classes are characterized by different splicing signals (Mount et al., 1992) or the presence of conserved elements (Pozzoli et al., 2007).

In this work, we develop a general statistical framework for indel dynamics and derive a set of models with increasing complexity that depict the length distribution of neutral sequences. We start with a simple model allowing indels of length one only, and reproduce TKF91 result stating that under a very weak deletion bias, arbitrarily large sequences are likely to appear. We extend this model by allowing indels of various lengths, and show that this allows the occurrence of neutral sequences even when the deletion bias is substantial. This is due to selection against deletions that encompass conserved regions at the neutral sequence borders, a phenomenon we term *border-induced selection*. Moreover, we suggest a model that includes small conserved elements embedded within the neutral sequence. The presence of these elements may significantly increase the neutral sequence length as they multiply the intensity of border-induced selection. Finally, we test how well our indel models explain the empirical intron length distribution in human. We show that the quantitative fit of the models improves with model complexity. Moreover, our framework provides an explanation for the multi-modality observed in the distribution of intron lengths.

## Results

### General model of length evolution

Our general goal is to understand how the length of neutral sequences evolves through generations. We start by describing a simple stochastic process for sequence length evolution. As we are only interested in length variation, substitutions are ignored, i.e., we implicitly assume that indel evolutionary dynamics is context independent, that is, the probability of indel events and their type does not vary as a result of substitutions. Further, we assume that the length can vary only due to indel events, and thus we ignore the possible contribution of rare events such as segmental duplications. In general, the variation of sequence length through generation can be described as follows:

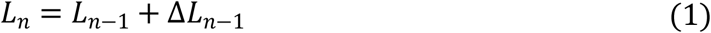

Where *L*_*n*_ is a random variable denoting the length of the sequence in generation *n*, and Δ*L*_*n*−1_ is a random variable that quantifies the sum of insertion and deletion lengths in the transition from generation *n* − 1 to generation *n*. Different assumptions regarding the indel dynamics would change the distributions of Δ*L*_*n*−1_, and thus the stationary distribution of *L*_*n*_. In the models proposed below, we focus on neutral segments that are bordered between highly conserved segments. We demonstrate the applicability of our models to introns, which we approximate as neutrally-evolving sequences.

### Human intron length distribution – empirical dataset for model validation

Below, increasingly complex models were tested for their fit to the human intron length distribution, as a representative of a large and well-curated empirical dataset. The human length distribution is characterized by the following features: mean intron length of ∼7,000 base-pairs (bp), standard deviation of ∼20,000 bp and a range that spans over five orders of magnitude: the minimal intron size is 30 bp, and the maximal is 1,160,411 bp (Piovesan et al., 2015). Furthermore, the distribution of the logarithm of the length is bimodal (Gotoh, 2018), with the main and minor modes at 2,100 and 100 bp, respectively (Figure 1).

**Fig. 1.**
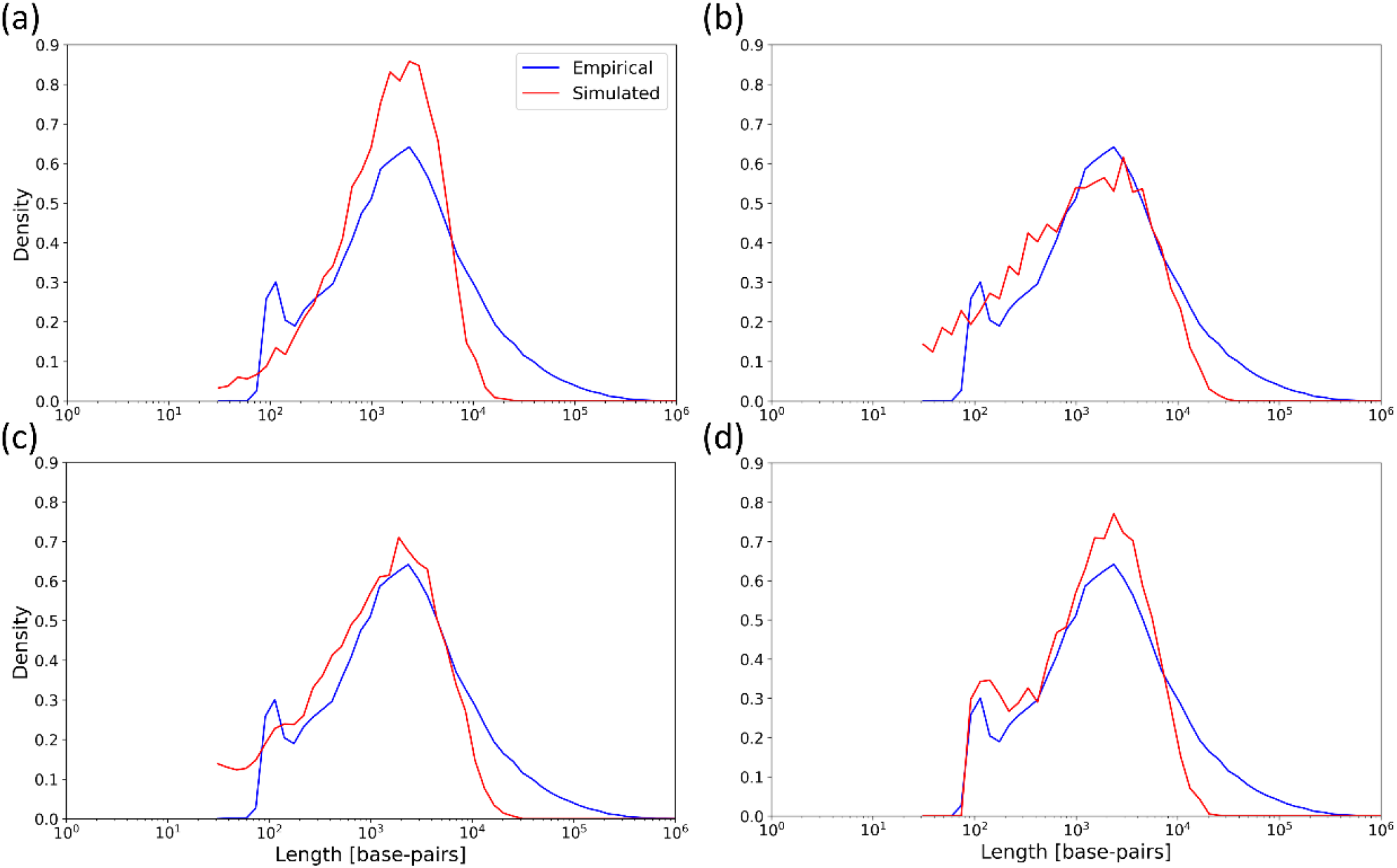
Empirical and simulated distributions of intron lengths in human. In each panel, the blue line shows a length distribution derived from the human intron empirical data. The red line is the distribution obtained using simulations with one of the models M1-M4. In all models, the length distribution was derived from 10,000 simulations. (a) The simulations are derived from M1, with the following parameters: *r* = 0.9995, *p*_*i*_ = 0.9995 · 10^−7^, *p*_*d*_ = 10^−7^. The MSE is 0.64; (b) The simulations are derived from M2 with the following parameters: *r* = 0.9975, *p*_*i*_ = 0.9975 · 10^−7^, *p*_*d*_ = 10^−7^, *μ*_*i*_ = 17, *μ*_*d*_ = 5. The MSE is 0.34; (c) The simulations are derived from M3 with the following free parameters: *r* = 0.983, *p*_*i*_ = 2.68 · 10^−8^, *p*_*d*_ = 10^−7^, *μ*_*i*_ = 16.5, *μ*_*d*_ = 4.5. The MSE is 0.31; (d) The simulations are derived from M4 that relies on the output of M3 model. The M3 parameters used here are: *r* = 0.9776, *p*_*i*_ = 2.65 · 10^−8^, *p*_*d*_ = 10^−7^, *μ*_*i*_ = 16.5, and *μ*_*d*_ = 4.5. The M4 model parameters are: *l*_*e*_ = 88, *l*_*i*_ = 35, and *p*_*c*_ = 0.69. The MSE is 0.29.

### Model with indels of size one

We start with a simple model (M1) that allows only insertions and deletions of size one and a uniform distribution of indel events along the sequence. We also assume that the sequence in question is placed between two conserved sequences that cannot be deleted. Therefore, even if the sequence length goes to zero in a certain generation, it can revive. This is analogue to the immortal link of the TKF91 model (Thorne et al., 1991). Under this model, the distribution of Δ*L*_*n*−1_ is:

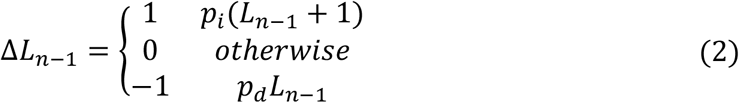

In this model, *p*_*i*_ and *p*_*d*_ are the probabilities of an insertion and deletion event, per character per generation, respectively. In each generation the length can vary by no more than a single character. We also assume that events are extremely rare, and thus, both *p*_*i*_(*L*_*n*−1_ + 1) and *p*_*d*_*L*_*n*−1_ are much smaller than 1.0, even for sequences longer than a million characters (Sung et al., 2016). Since insertions occur between characters, there is an additional place for insertions compared to deletions, i.e., deletions can only occur upstream to each character while an insertion can also occur downstream to the last character. For example, if an intron is of length three bases, insertions can occur at four possible locations, while deletions can occur at only three locations, i.e., upstream of each base.

Given the stochastic process described above, taking expectations from both sides of equation (1) yields:

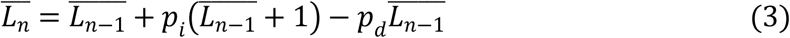

Equation (3) coincides with the model of He et al. (2019). The solution for equation (3) for the case in which *p*_*i*_ = *p*_*d*_ is a linear growth, where *L*_0_ is the expectation of the sequence length at the beginning of the process:

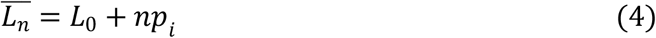

When *p*_*i*_ ≠ *p*_*d*_ the solution is:

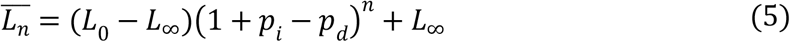

where 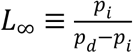. This notation is used as the length converges to *L*_∞_ when *p*_*i*_ < *p*_*d*_, and not to *p*_*i*_ as reported by He et al. (2019). If *p*_*i*_ > *p*_*d*_, the exponential term grows to infinity. Figure 2 demonstrates the behavior of the solution of equation (3) for the three regimes: *p*_*i*_ > *p*_*d*_, *p*_*i*_ = *p*_*d*_ and *p*_*i*_ < *p*_*d*_.

**Fig. 2.**
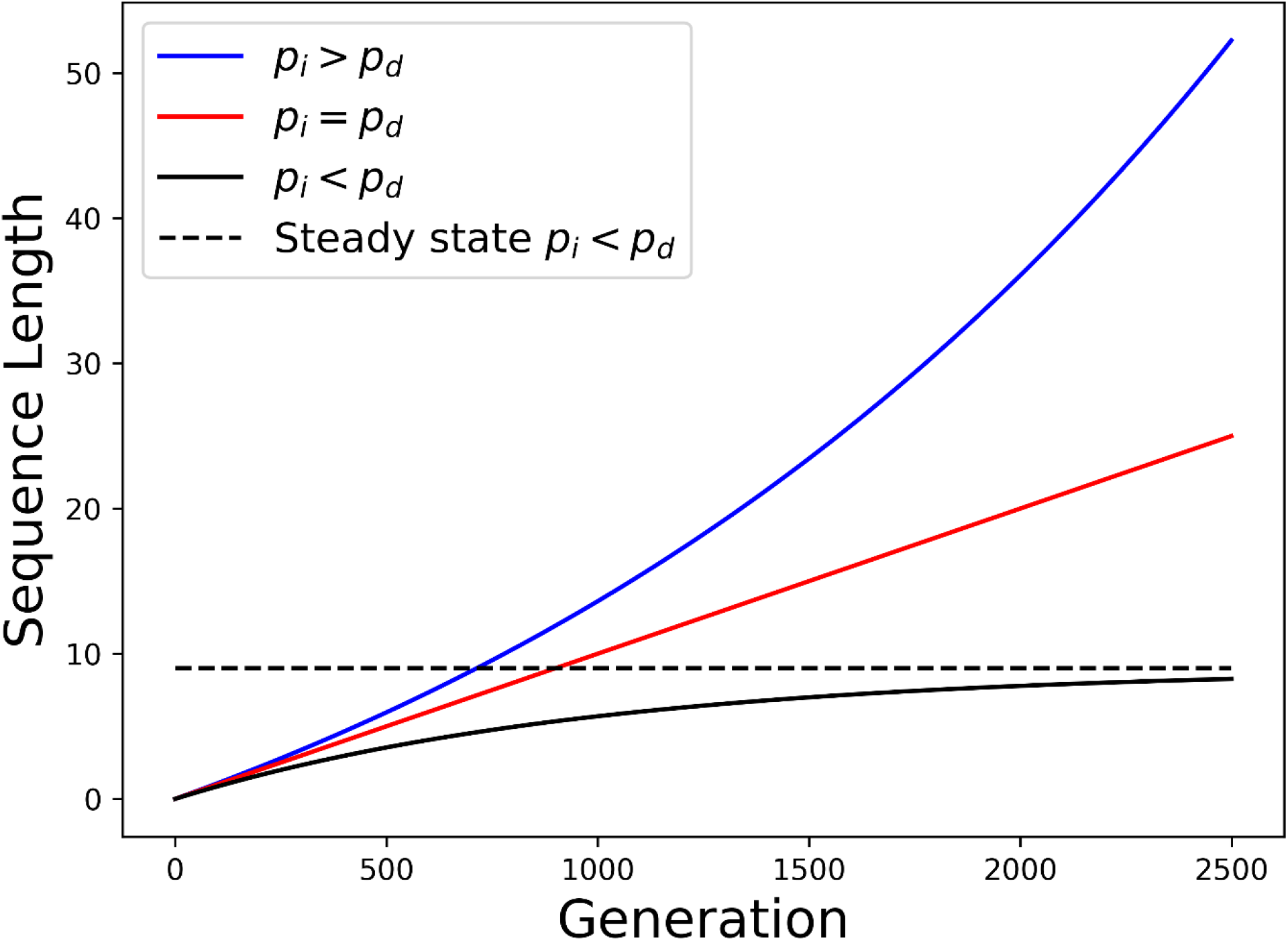
The expectation of the model with single-character indels (M1). Three types of solution are possible: exponential growth when *p*_*i*_ > *p*_*d*_, linear growth when *p*_*i*_ = *p*_*d*_, exponential decay to a steady state when *p*_*i*_ < *p*_*d*_. The parameters used to generate the graphs were: *L*_0_ = 0, *p*_*d*_ = 0.01, *p*_*i*_ = (0.0105 *or* 0.01 *or* 0.009).

We will focus on the deletion bias regime, i.e., *p*_*i*_ < *p*_*d*_, as it was repeatedly reported that deletions are more common than insertions. The steady state length, *L*_∞_, depends solely on the ratio between the insertion and deletion probabilities, 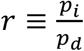 (multiplying the value of *p*_*i*_ and *p*_*d*_ by a fixed factor has no effect on the stationary distribution – it only affects the time till convergence, see Appendix 1):

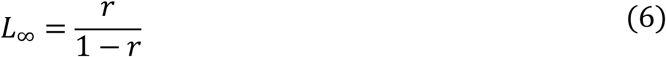

Equation (6) shows that *L*_∞_ can be arbitrarily long by selecting the appropriate *r* value. For example, when *r* = 0.9, 0.99, 0.999 then *L*_∞_ ≈ 9, 99, 999, respectively. Under this model, when the sequence is shorter than *L*_∞_, it has an insertion bias even though *p*_*i*_ < *p*_*d*_. For example, when the sequence length is one, there are two possible insertions and a single possible deletion, thus if 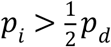 the sequence will have an insertion bias. Of note, equation (6) is the same as reported in the TKF91 model (Thorne et al., 1991).

The above model clearly does not fit the human empirical intron data. Shown in figure 1 is the distribution of the intron lengths of the human genome (see Material and Methods). The mode of this distribution is 2,100 bases. Simulations with the above model allow obtaining estimate of the stationary distribution for each value of *r*. We searched for the value of *r* that provides the best fit in terms of mean squared error (MSE) between the empirical and simulated length distributions of human introns (see Material and Methods). The optimal value of *r* was 0.9995 with an MSE of 0.64. Although for this value of *r* the main mode of the empirical distribution matches the mode of the simulated stationary distribution, the two distributions vary greatly with respect to their shapes (Figure 1a). Specifically, while the empirical distribution has a heavy right tail, these long introns are missing from the stationary distribution generated by the model. In addition, the empirical distribution has a second mode near 100 bp, which is missing from the simulated distribution. Given this discrepancy, we now turn to a more complex model that relaxes the over-simplified assumption that all indels are of size one.

### Model with indels of fixed arbitrary size

We generalize the above-described model by adding parameters *μ*_*i*_ and *μ*_*d*_ that are the insertion and deletion lengths, respectively. Of note, these lengths are considered constant (below, we relaxed this assumption by allowing a distribution of indel sizes). Under M2 the distribution of *L*_*n*−1_ is:

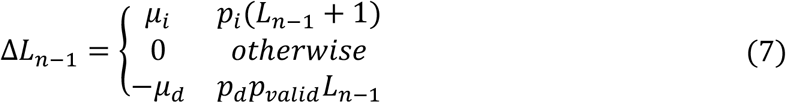

We note that because deletions are no longer restricted to have a length of one, some deletions may extend from the neutral sequence to its conserved flanking regions, entailing substantial fitness reduction, and thus such deletions are rejected. This is reflected in the extra factor *p*_*valid*_ in equation (7). Given the value of *L*_*n*−1_ and *μ*_*d*_, *p*_*valid*_ can be computed by:

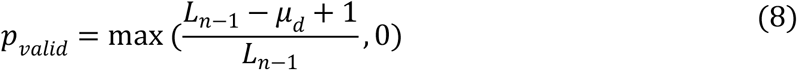

Of note, if the proposed deletion length is larger than the current sequence length, equation (8) will assign a probability of zero to *p*_*valid*_, suggesting that neutral sequence segments are immune to deletions larger than their size. Under this scenario, there is a bias for insertions in neutral sequences that are very short (See also Ptak and Petrov, 2002).

Taking the expectation of both sides of equation (1), accounting for the distribution of Δ*L*_*n*−1_ as in equations (7-8) yields:

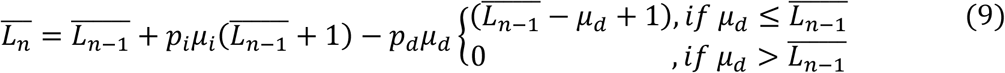

The first two terms on the right-hand side resemble equation (3), just that the second term is multiplied by the insertion length *μ*_*i*_. The third term indicates that no deletions are allowed when 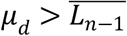. As expected, when we choose *μ*_*i*_ = *μ*_*d*_ = 1, equation (9) reduces to equation (3). Of note, equation (9) is effectively a three parameters difference equation. Let *r* be the ratio between the expectation of the insertion length and the expectation of the deletion length: 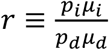. Note that *r*, as defined for M1 (equation 6), is a special case of the *r* in M2, when *μ*_*i*_ = *μ*_*d*_ = 1. Using these definitions, we can rewrite equation (9) with three parameters *r, p*_*d*_, and *μ*_*d*_:

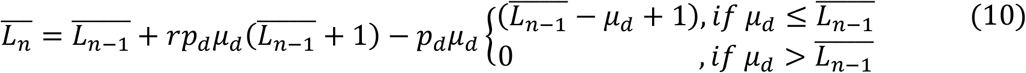

When the steady state length, *L*_∞_, is substantially larger than *μ*_*d*_, we can approximately ignore the 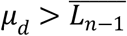 condition and solve the following equation:

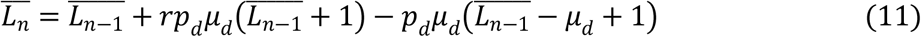

The solution of equation (11) resembles the solution of equation (3):

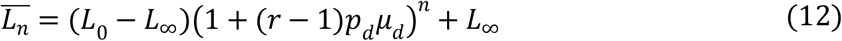

The steady state sequence length, *L*_∞_ is:

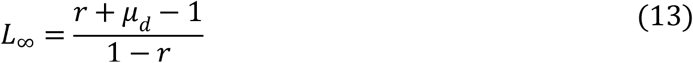

It is interesting to compare the properties of M2 and M1. First, as expected, if *μ*_*d*_ and *μ*_*i*_ are set to be 1, M2 reduces to M1. Second, when *r* is close to 1, *L*_∞_ under M2 is roughly *μ*_*d*_ fold larger than *L*_∞_ under M1. Third, when *r* is close to 0, there are substantially more deletions than insertions, and thus the 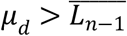 condition of equation (10) may not be negligible. Under such a high deletion regime, many deletions are rejected and *L*_∞_ should be larger than the value predicted by equation (13). Hence, in this case, the value in equation (13) can be considered as a lower bound for the steady-state length.

To fit this model to the human empirical intron data, we assume that the mean insertion and deletion lengths are 16.5 and 4.5 base-pairs, respectively, as reported by Matthee et al. (2007) for introns in mammals. We use 17 and 5 base-pairs, as this model supports only integers. We scanned the *r* parameter, using the same procedure we applied for M1, and the *r* value that yielded the optimal fit was 0.9975 with an MSE of 0.34, which is a substantial improvement over the MSE obtained for M1. The inferred *r* value is slightly lower than that obtained using the M1 model. The increased fit and the fact that the shape of the M1 and M2 distributions are different emphasize the importance of the conserved regions at the edges of the neutrally evolving sequence, i.e., boundary-induced selection. Figure 1b shows that despite the increased fit as measured by the MSE value, substantial discrepancies remain between the simulated and the empirical distributions.

### Model with indels of varying sizes

We generalize the above model by relaxing the assumption that the insertions and deletions are of constant length. Thus, in M3 we assume that the length of each indel is drawn from a specified distribution. Let *f*(·) and *g*(·) be the length distributions of insertions and deletions, respectively. The distribution of *L*_*n*−1_ under this model is:

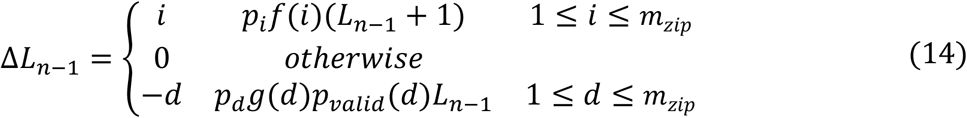

The probability that a deletion is valid depends on the deletion length, as described in equation (8). The longer the deletion length *d* is, the smaller *p*_*valid*_(*d*) is. In equation (14), *p*_*valid*_(*d*) is a function of *L*_*n*−1_, which complicates the analytic computation of the expectation of Δ*L*_*n*−1_.

In this work, we assume a truncated Zipfian distributions for both insertions and deletions, as in Loewenthal et al. (2021):

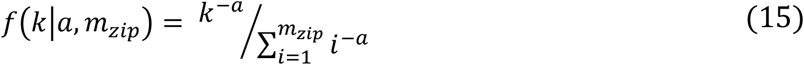

This distribution has two parameters *a* and *m*_*zip*_, which control the shape of the distribution and the maximally allowed indel length, respectively. We assume that both insertion and deletion lengths are Zipfian distributed, but we allow different *a* parameters for insertions and deletions. Unless otherwise stated, *m*_*zip*_ is set to 150 throughout this work. We denote by *μ*_*i*_ and *μ*_*d*_ the expectations of the truncated Zipfian distribution for insertions and deletions, respectively.

In Appendix 2, we show, using simulations, that the mean value of this distribution, *L*_∞_, for M3 is about 4.5-fold higher compared to model M2 when using the same set of parameter values. The higher mean in M3 compared to M2 stems from the higher probabilities that proposed long deletions are rejected.

The mean of the stationary distribution in M3 (i.e., *L*_∞_) is often larger than the mean deletion length, even under a very strong deletion bias regime. For example, when *r* is 0.25, *μ*_*d*_ = 15, and *μ*_*i*_ = 5, the mean length is 23.2 bp (Figure 3). This can be explained by the fact that when the segment length is shorter than the mean deletion length, most deletions are rejected, and thus, effectively, a strong bias for insertions exists.

**Fig. 3.**
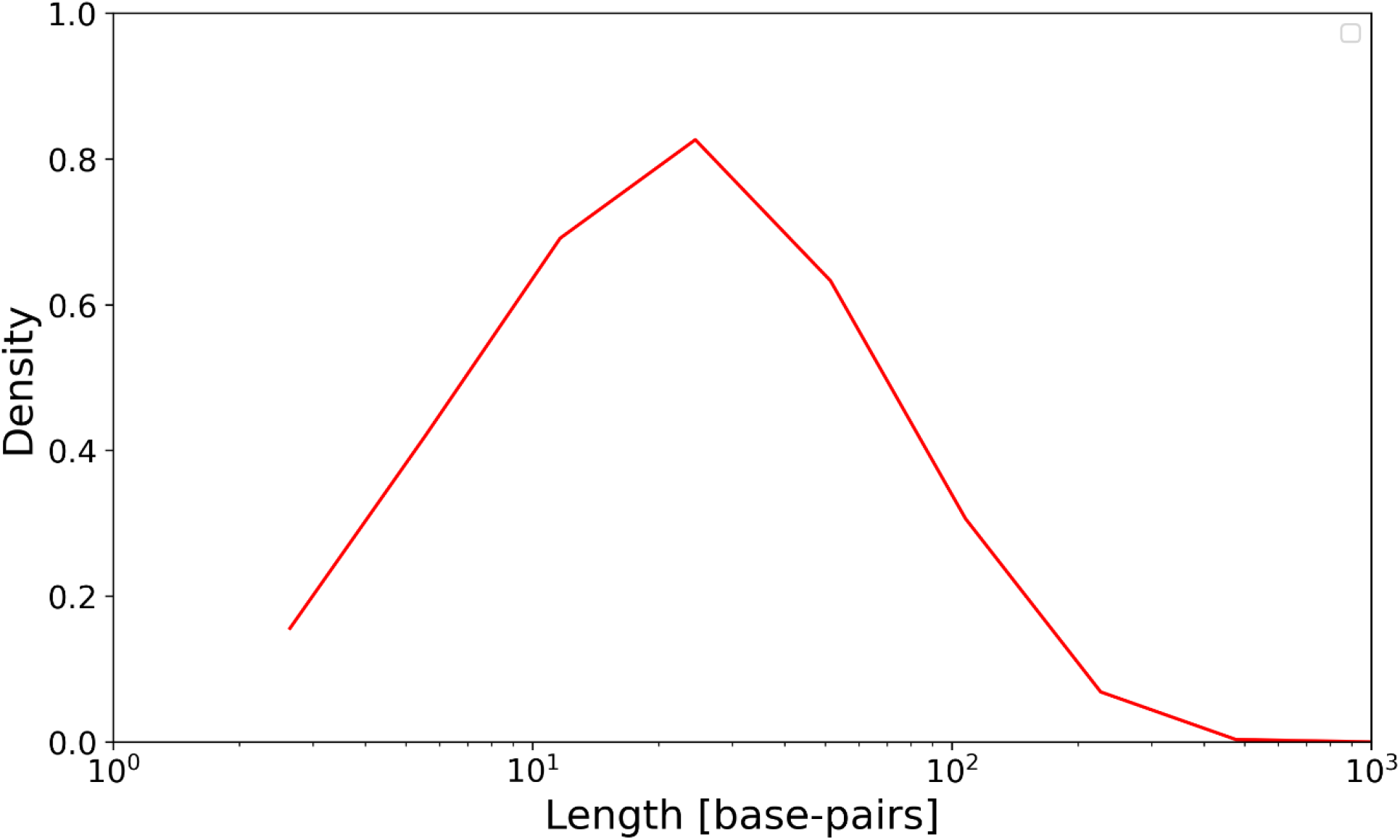
Long neutral sequences are probable in M3 under a high deletion bias regime. The red line is the distribution obtained using M3 under a strong deletion bias regime (*r* = 0.25). The length distribution was derived from 10,000 simulations with the following parameters: *p*_*i*_ = 7.5 · 10^−9^, *p*_*d*_ = 10^−8^, *μ*_*i*_ = 5, *μ*_*d*_ = 15.

Applying this model to the human intron length distribution, we found that the best fit is obtained with *r* = 0.983, yielding an MSE of 0.31. In this computation we applied the mean indel lengths as reported in Matthee et al. (2007), i.e., *μ*_*d*_ = 4.5 and *μ*_*i*_ = 16.5 (Figure 1c). The modes of the empirical and simulated distributions are similar, however, major discrepancies between the shapes of the two distributions exist: the distributions mean and standard deviations are (6,793; 21,860) and (1,684; 2,354) bp for the empirical and simulated distributions, respectively. There is also an additional peak in the empirical distribution (bimodality) that is absent in the simulated distribution.

### Conserved segments

We propose a toy statistical model, M4, to qualitatively demonstrate that conserved segments embedded within the neutrally evolving sequence may explain the gap between the theoretical and empirical distributions. Accordingly, in M4 we assume that in each neutral sequence there is some probability, *p*_*c*_, that it includes a single conserved sequence of length *l*_*i*_ within it. Let *l*_*e*_ denote the total length of the conserved sequences in the edges of an intron (i.e., the 5’ and the 3’ splice sites). To simulate this model, for each neutral sequence, a Bernoulli trial is executed with probability *p*_*c*_ to decide if there is a conserved sequence within the intron sequence (in M4, we only allow a single internal conserved sequence). If there is no conserved sequence, then the length of the intron is the sum of *l*_*e*_ and the length of a single stochastic simulation under M3. If a conserved sequence is introduced, the length of the intron is the sum of *l*_*e*_, *l*_*i*_, and the lengths of two stochastic simulations under M3. For simplicity, we assume that the parameters *l*_*e*_, *l*_*i*_, *p*_*c*_ are the same for all the simulated introns. The optimized simulated distribution is shown in figure 1d. In contrast to the fit in model M3, now the simulated distribution is bimodal similarly to the empirical distribution. The fit between the two distributions slightly improved (the MSE decreased from 0.31 to 0.29), the value of *r* decreased to 0.9776, and the mean and the standard deviation of the sequence length both slightly increased, reaching (2,149; 2,365) bp. The fitted parameters under M4 are *l*_*e*_ = 88, *l*_*i*_ = 35, and *p*_*c*_ = 0.69. The high *p*_*c*_ value suggests that most of the introns have a conserved internal segment. Thus, conserved segments may explain the low peak in the intron length distribution, it widens the length distribution, and resulted in a lower value of *r*.

Here and in previous works (Gotoh, 2018), the intron length values are transformed using the log function prior to their visualization as a distribution. This is justified, as the intron lengths are spread over five orders of magnitude. This result in a bimodal distribution. Of note, when the same empirical distribution is plotted without log scaling, the bimodality disappears (Figure 4a). Our analyses suggest that inclusion of conserved segments (both within and in the border of introns) led to the appearance of bimodality in the log-scale and to longer introns. Specifically, it is the introduction of conserved elements in the borders of introns that mostly explains the bimodality, and the presence of internal conserved elements that lead to the generation of longer introns (Figure 4b).

**Fig. 4.**
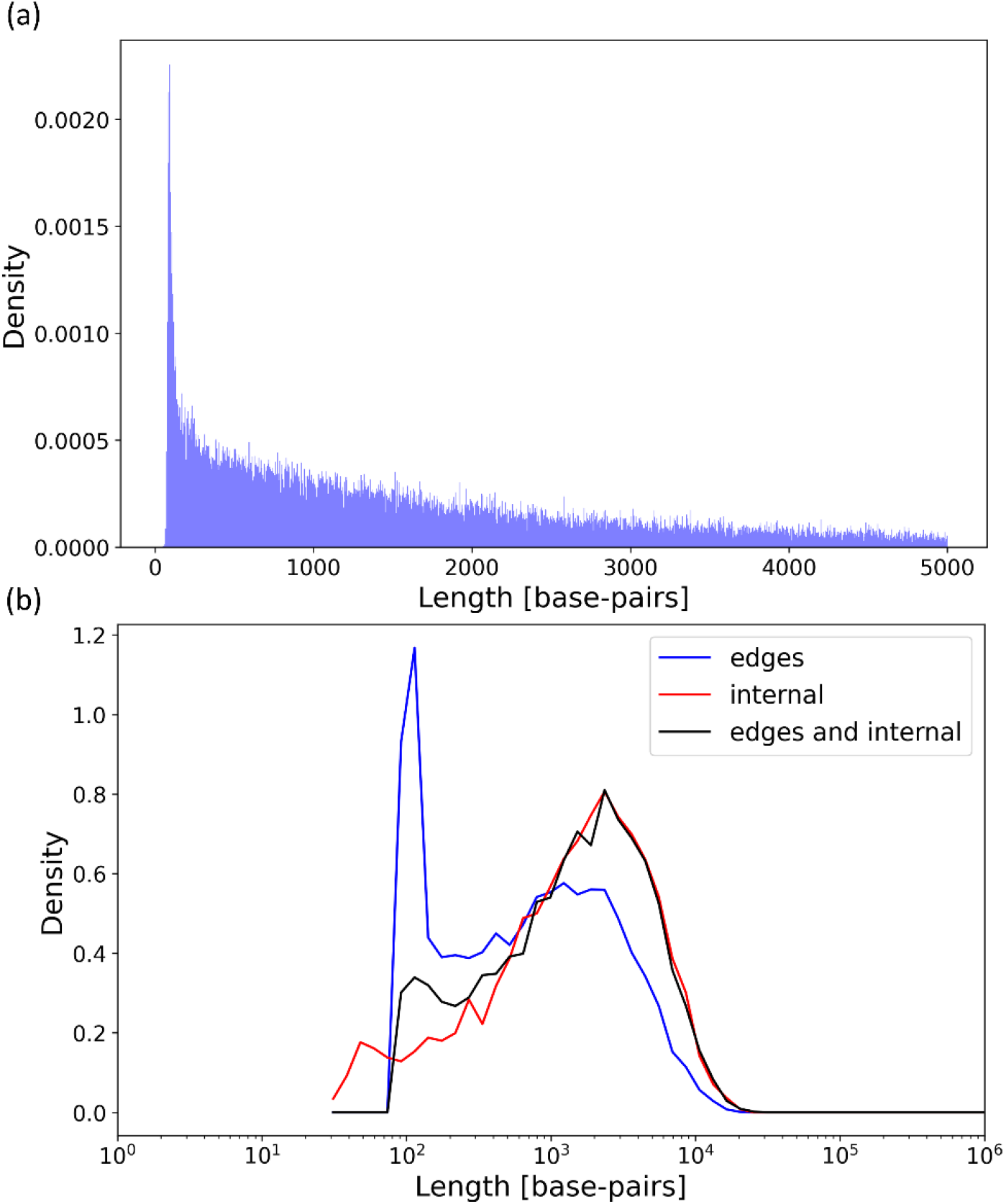
Empirical length distribution in linear scale and M4 parameter. (a) The human intron length distribution of introns shorter than 5,000 bp in a linear scale. There is a single mode, as opposed to the logarithmic scale. Thus, the two modes are an artifact of the logarithmic transformation. (b) Three M4 simulations with ran with the same M3 parameters: *r* = 0.9776, *p*_*i*_ = 2.65 · 10^−8^, *p*_*d*_ = 10^−7^, *μ*_*i*_ = 16.5, and *μ*_*d*_ = 4.5. In blue, we use the M4 parameters: *l*_*e*_ = 88, *l*_*i*_ = 0, and *p*_*c*_ = 0. Thus, we only add a constant to the M3 simulation, reflecting the conserved intronic splice sites at the edges of the intron. It can be seen that bimodality emerged. The mean length is 1,295 bp. In red, we use the M4 parameters: *l*_*e*_ = 0, *l*_*i*_ = 35, and *p*_*c*_ = 0.69. These parameters dictate the presence of introns with an internal conserved segment, but there are no conserved segments at the edges of the introns. The mean value increased to 2,067 bp, reflecting how internal conserved segments may significantly increase the intron lengths. In black, running the simulations with conserved segments both within and at the edges of the intron. The M4 parameters were *l*_*e*_ = 88, *l*_*i*_ = 35, and *p*_*c*_ = 0.69. The mean length is 2,158. Of note, this distribution was generated with the same parameters as in figure 1d and the small differences reflect stochastic variations.

## Discussion

In this work, we presented several increasingly complex models for the length distribution of neutrally evolving sequences. Critical to our models is the assumption that neutrally evolving segments are placed between highly conserved sequences. We focused on the deletion bias regime, which was shown in a large number of studies to be prevalent across all domains of life (Fitch, 1973; Graur et al., 1989; De Jong and Rydén, 1981; Kuo and Ochman, 2009; Loewenthal et al., 2021; Mira et al., 2001; Ogata et al., 1996; Van Passel et al., 2007; Tao et al., 2008; Zhang and Gerstein, 2003). It was previously suggested that this deletion bias leads to shrinkage of genomes over evolutionary times (Petrov, 2002). Here, we showed that the placement of conserved flanking sequences can lead to the emergence of long sequences, even in a high deletion bias regime. The counter-intuitive result that long neutrally evolving sequences can emerge even under a strong deletion bias is due to the rejection of deletions that invade the highly conserved borders of the neutral sequences. We hence propose the term border-induced selection for this phenomenon.

To test the fit of our models to empirical genomic data we studied the length distribution of human introns, which are thought to evolve mostly neutrally (Resch et al., 2007) and are in between exons, which are generally highly conserved (Siepel and Haussler, 2004). Using the M3 model, we reconstructed the main mode of the empirical distribution. However, M3 does not reproduce a secondary lower peak of the distribution and does not explain the extremely high variance of the lengths of introns, which in this case spans over five orders of magnitude. Yet, the M3 model does not account for conserved segments within and at the edges of introns. Examples for such conserved segments are the 3’ and 5’ splice sites, as well as intron splicing enhancers and silencers (Barash et al., 2010; Cooper, 2010; Lin et al., 2016). We modeled the presence of conserved segments within introns using M4, and it resulted in both the emergence of a second peak and a slight increase in the variance. Of note, M4 only allows a single intermediate conserved segment, and we expect that a more elaborate model that allows multiple conserved internal segments will better explain the presence of very long introns.

As is often the case with models, many assumptions are clear over-simplifications of biological realism. First, the output of the stochastic model depends on the indels length distribution. As in previous work, we assumed that this distribution follows a truncated Zipfian distribution (Loewenthal et al., 2021) with a cut-off of 150 characters. In our work, we did not study if this is the best fitting distribution, and it is possible that other distributions may provide better fit to the data. Our model also assumes perfect neutrality of the sequence of interest and perfect conservation of the bordering conserved elements. This is also true for the conserved regions within introns. The effect of relaxing these assumptions needs to be further studied. Of note, M4 is not a genuine stochastic model with specified parameters controlling the probability of emergence and loss of conserved regions However, we expect that a more complex model, which addresses these limitations, will not change the main result of our model, namely, that neutral sequences are not purged under a deletion bias regime. Moreover, mobile genetic elements, microsatellites, and genome rearrangement events are all ignored in our study. Clearly, these factors should be integrated when moving towards complex models that aim to capture the main forces dictating genome dynamics evolution. Finally, throughout this work, we assumed that the empirical length distributions are in equilibrium and we thus compare them to the stationary distributions of our models. It may be the case that this assumption too is an oversimplification of reality. While in this work we focused on the evolution of intron lengths, our models provide a framework to study length distribution of other neutrally evolving sequences such as prokaryotic spacers (Rédei, 2008).

Our analyses show that as we move to increasingly more complex models, the insertion-to-deletion rate ratio, *r*, gets further away from the value of one. Equation (13) indicate that as *r* gets closer to one small perturbations of *r* lead to sharp changes in intron length distribution. Since in our models *r* is closer to one, the mode is unstable. For example, changing the value of *r* between values such as, say, 0.9995 and 0.9998, would generate distributions with very different means: from 2,000 to 5,000 bp in M1, respectively. Indeed, the decrease in *r* as we move to more advanced models, reaching an *r* of 0.98 in M4 lends an additional level of justification for these advanced models. We anticipate that incorporating multiple conserved elements will further lead to more stable models.

Previous studies provide indirect support for our proposed models. First, Pozzoli et. al (2007) compared mouse and human introns, and showed that the deletion rate is higher for long introns, in line with our models, because deletions in short introns are often rejected, while in long introns there is little to none border-induced selection. Pozzoli et. al also examined introns of similar length, and found that the number of conserved sequences is negatively correlated to deletion rate, again in line with the existence of border-induced selection. Moreover, the authors also showed that almost all introns longer than 10,000 bp harbor conserved sequences, emphasizing the important role conserved segments play in generating the heavy tail of the intron length distribution. Second, Yang et al. (2021) have recently showed that within a genome, the intron size is correlated to the alternative splicing level and prevalence. Sironi et al. (2005) showed a correlation between the logarithm of intron length and the number of conserved sequences within the intron. These observations can fit a general model, in which tight regulation of splicing is associated with conserved intronic regulatory elements, which, as we showed, lead to long introns. Third, it was shown that first introns are much longer, typically about double, than other introns, which may be partially explained by the observation that functional motifs are more frequent in first introns (Bradnam and Korf, 2008). This observation further supports the M4 model, in which the presence of conserved segments leads to longer introns.

Our model provides a plausible explanation for the extremely large variance in intron lengths within a species. However, it does not directly explain differences in distributions among species. One trivial explanation is that the model parameters themselves, evolve. Thus, different species have different insertion to deletion rate ratios, and possibly, different propensity for the emergence of conserved regions within introns. These factors may be relevant not only to the distribution of intron lengths, but rather, for the entire genome size. Indeed, eukaryotes generally have a lower deletion bias than prokaryotes (Kuo and Ochman, 2009), which may partially explain the higher eukaryotes genome sizes and their higher variation (Bohlin and Pettersson, 2019). It was previously shown that the total indel rate is negatively correlated to the effective population size (Sung et al., 2016). It was also shown that the effective population size times the mutation rate is correlated to the introns mean length (Lynch and Conery, 2003). A dependence between the insertion-to-deletion ratio and the effective population size, if exists, may help explain this relationship: smaller population size leads to an increased *r*, which in turn leads to longer introns.

## Materials and Methods

### Intron length distribution

We downloaded the canonical genome of human from the UCSC Genome Browser database (Rosenbloom et al., 2015). The canonical genome introns annotation is based on the longest coding sequence isoform for each gene. The complete distribution of intron lengths in the canonical human genome is provided at https://github.com/elyawy/Luigi (last accessed June 29th, 2022).

### Simulations and optimization of model parameters

The simulations of M1, M2, and M3 are based on the Gillespie algorithm (Gillespie, 1977). We used discrete generations, and thus waiting times were geometrically distributed. The number of generations needed to reach stationarity is dictated by the transient part of equation (12), i.e., (1 + (*r* − 1)*p*_*d*_ *μ* _*d*_)^*n*^. We simulated until this factor was below 10^−6^.

Model parameters for M1-M3 were optimized using a grid search over the *r* parameter in the range [0.37, 0.9999]. The optimal *r* parameter had the lowest mean square error between the simulated and empirical length distribution (in logarithmic scale). For M4, we heuristically searched for the values of (*r, l*_*e*_, *l*_*i*_, and *p*_*c*_) that best fit the empirical distribution according to the mean square error criterion. This was done using the module optimize of Python SciPy package (Virtanen et al., 2020) using the ‘trf’ option, which is based on the trust region algorithm described in Gould et al. (1999).

### Source code and implementation details

The source code and documentation of the C++ (models M1-M3) and Python (model M4) implementation of the stochastic simulations are available at https://github.com/elyawy/Luigi (last accessed June 29th, 2022).

## Acknowledgments

G.L., E.W., N.N, and L.G were supported in part by a fellowship from the Edmond J. Safra Center for Bioinformatics at Tel Aviv University. We thank Jotun Hein and Alon Itzkovitch for fruitful discussion.

## Appendix 1

**Table A1.** Multiplying the value of *p*_*i*_ and *p*_*d*_ by a fixed factor has no effect on the stationary distribution. Each row in the table represents a set of 10,000 simulations under M1 with the parameters *p*_*i*_ and *p*_*d*_ specified in the first two columns. Each set of simulations were run until convergence to the stationary distribution. In each row, the parameters *p*_*i*_ and *p*_*d*_ were multiplied by 0.1 compared to the row above, so the *r* is fixed to 0.99 in all rows. The empirical means and standard deviations (sd) of the various sets are similar. We also performed a two-sided Kolmogorov–Smirnov (KS) test between the first simulation set and the other sets. The null hypothesis, namely that the distributions are the same, cannot be rejected.

| $p_i$ | $p_d$ | $r$ | Empirical<br>mean | Empirical<br>sd. | KS. Stat. | KS. P value |
| --- | --- | --- | --- | --- | --- | --- |
| 9.90E-07 | 1.00E-06 | 0.99 | 98.50 | 99.86 | - | - |
| 9.90E-08 | 1.00E-07 | 0.99 | 99.03 | 98.84 | 0.01 | 0.34 |
| 9.90E-09 | 1.00E-08 | 0.99 | 98.23 | 99.22 | 0.01 | 0.63 |
| 9.90E-10 | 1.00E-09 | 0.99 | 100.24 | 102.71 | 0.01 | 0.58 |

## Appendix 2

We computed *L*_∞_ for M3 using simulations with various values for *r, μ*_*d*_, and *μ*_*i*_. Here, *μ*_*i*_, and *μ*_*d*_ denote the expectations of the truncated Zipfian distribution for insertions and deletions, respectively. Our simulations suggest that a strong linear correlation exists between the simulated value of *L*_∞_ and the *L*_∞_ values calculated based on equation (13). Let *κ*_*zipf*_ be the slope of the regression line (assuming an intercept of zero). Thus,

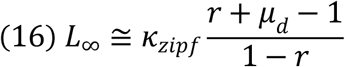

Note that in this case, the value of *r* depends on both *μ*_*d*_ and *μ*_*i*_. We also note that the truncated Zipfian distribution depends on two parameters: *a* and *m*_*zip*_. The values of *μ*_*d*_ and *μ*_*i*_ can be computed given these two parameters. In equation (16), *κ*_*zipf*_ may vary depending on *m*_*zip*_. The correlation coefficient between the estimated *L*_∞_ based on equation (13) and *L*_∞_ estimated using simulations was found to be higher than 0.97 for all tested *m*_*zip*_ values (figure S1).

The fact that the slope of the regression line is higher than one in all cases (Figure S1), suggests that introducing variation to the indel lengths pushes the distribution of sequence lengths to higher values, including increasing the average length *L*_∞_ by a factor *κ*_*zipf*_>1. We hypothesize that the reason for the shift in sequence lengths is due to a reduction of the expectation of the mean length of accepted proposed deletions, which stems from the variation of indel lengths. Indeed, the expectation of the length of accepted proposed deletions for an arbitrary deletion length distribution *g*(*d*) for a sequence of length *L, E*[*a. d*], is given by *E*[*a. d*] = ∑_*i*_ *ig*(*d*) *p*_*valid*_. This expectation has a compact form when the maximal deletion length is lower than 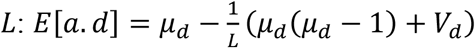 where *μ*_*d*_ and *V*_*d*_ are the mean length and variance of *g*(*d*), respectively. As expected, when *L* → ∞, only the first term contributes so *E*[*a. d*] = *μ*_*d*_, but for a finite *L*, the negative second term reduces *E*[*a. d*], and thus the reduction is higher when variation in the allowed deletion length is introduced.

