## Supplementary material for "The evolutionary dynamics that retain long neutral genomic sequences in face of indel deletion bias: a model and its application to human introns": figure S1

**Fig. S1.**

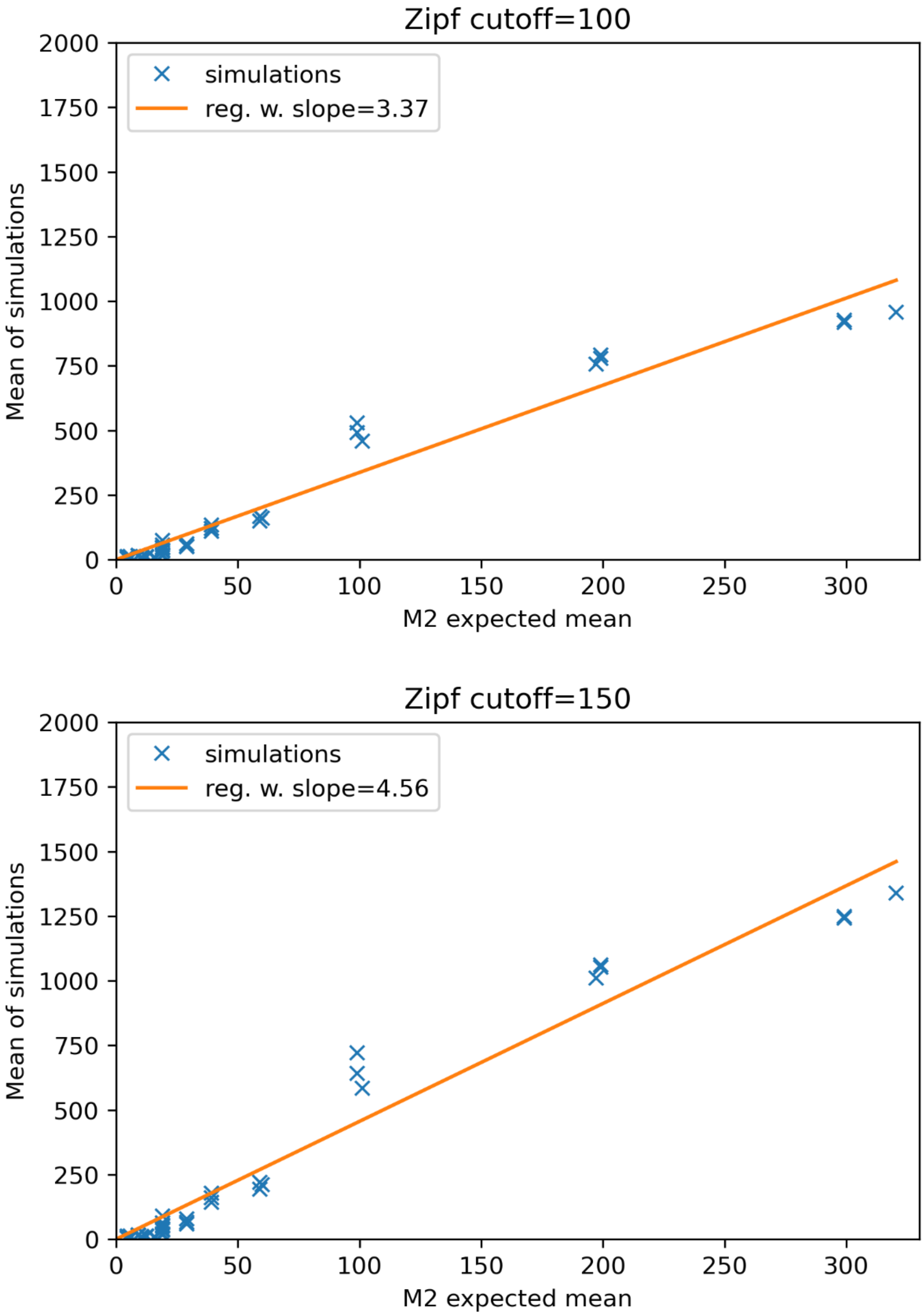
Zipf cutoff=100
simulations
reg. w. slope=3.37
Mean of simulations
M2 expected mean
2000
1750
1500
1250
1000
750
500
250
0
0
50
100
150
200
250
300
Zipf cutoff=150
simulations
reg. w. slope=4.56
Mean of simulations
M2 expected mean

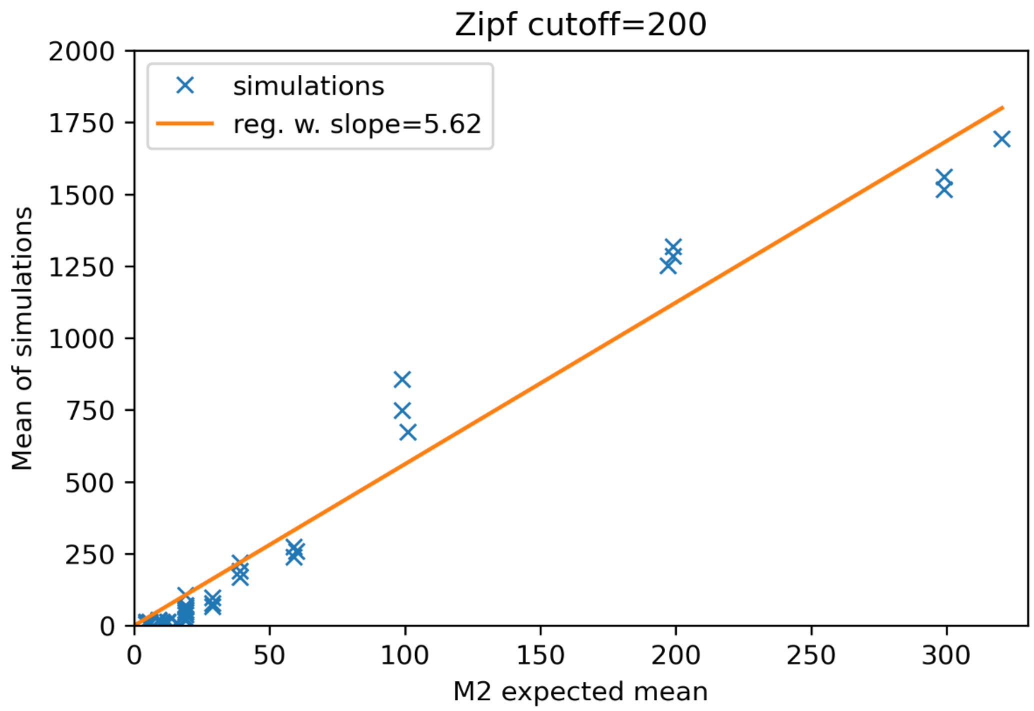


**The means of simulations under M3 are highly correlated to those under M2.** The slope of the regression line depends on the indel length distribution variance. The figure is composed of three sub figures for M3 simulations preformed with three cutoffs of the truncated Zipfian distribution: 100, 150, 200. Each blue 'x' symbol in a subfigure corresponds to a set of 10,000 simulations with the same M3 parameters. The x-axis is the M2 expected mean of these parameters according to equation (13), and the y-axis is the mean of the simulations set. Each sub-figure is composed of 45 sets of simulations with the following parameters combinations: $\mu_i = 5, 10, 15$, $\mu_d = 5, 10, 15$, $r = 0.1, 0.25, 0.5, 0.75, 0.95$, $p_d = 10^{-8}$, and $p_i = rp_d\frac{\mu_i}{\mu_d}$ (three options of $\mu_i$, three options of $\mu_d$, and five options of $r$, therefore 45 simulations). The orange line in each subfigure is a regression with zero intercept line of the 'x' symbols data. The regression slope in the subfigures is 3.37, 4.56, and 5.62 for the truncated Zipfian cutoffs 100, 150, and 200, respectively. Of note, that the regression slope increases as the Zipfian cutoffs increases because for a given mean, the variance of truncated Zipfian distribution increases when the cutoff increases.
